# A newly developed model of atherosclerosis caused by the stress of metabolic syndrome in apolipoprotein E knockout mouse

**DOI:** 10.1101/2025.03.24.645136

**Authors:** Riku Tsudome, Yasunori Suematsu, Kohei Tashiro, Hidetaka Morita, Akihito Ideishi, Shin-ichiro Miura

## Abstract

**Background:** Atherosclerotic disease is influenced by multiple factors including hypertension, dyslipidemia, and diabetes and it is rarely due to a single pathology in clinical conditions. In this study, we created a mouse model of metabolic syndrome and aimed to elucidate the mechanism of atherosclerosis in metabolic syndrome.

**Methods:** Twelve-week-old male apolipoprotein E knockout mice were fed a high-fat diet to produce dyslipidemia, administered angiotensin II continuously to develop hypertension, and infused with streptozotocin intraperitoneally at 18 weeks to produce diabetes mellitus for metabolic syndrome (METS) mice. Control (CTL), high-fat diet alone, angiotensin II alone, and streptozotocin alone mice were also produced and the results in the various groups were compared.

**Results:** The METS group showed increased body weight, blood pressure, and serum total cholesterol. The METS group exhibited a significantly higher rate of atherosclerosis formation than the CTL group (METS; 25.7±11.6 %, CTL: 3.3±1.5 %, p<0.001). The expressions of nicotinamide adenine dinucleotide phosphate oxidase 2, nuclear factor-kappa B1, and matrix metalloproteinase-2 in the METS group were significantly higher than those in the CTL group by reverse transcription-polymerase chain reaction analysis.

**Conclusions:** The METS model mouse showed severe atherosclerosis caused by oxidative stress, inflammation, and atherosclerotic plaque formation. This model is expected to be useful as a model that more closely resembles the pathophysiology of clinical atherosclerosis.

**Perspective:** *What Is New?:* This study introduces a novel metabolic syndrome mouse model that closely resembles human atherosclerosis by integrating hypertension, dyslipidemia, and diabetes into an apolipoprotein E knockout mouse.

*What Are the Clinical Implications?:* The metabolic syndrome mouse model demonstrates severe atherosclerosis driven by oxidative stress and inflammation, making it a valuable tool for testing therapeutic strategies targeting metabolic syndrome-related cardiovascular diseases.

## 3. Introduction

Atherosclerosis is a highly complex inflammatory process. Repeatedly evaluating for the progression of vascular inflammation in clinical studies is challenging and this places significant emphasis on preclinical experiments. In the field of atherosclerosis, animal experiments began in 1908 when Alexander Ignatowski established the first animal model using rabbits that had been fed a cholesterol-rich diet to study atheromatous plaque formation. Since then, studies on atherosclerosis have primarily used rabbits [1]. Studies on atherosclerosis model mice began in the latter half of the 20^th^ century, and to this day, mice remain the primary species for research due to their reproductive capacity, ease of genetic manipulation, and the ability to form atherosclerosis in a relatively short period [2, 3]. However, mice have a different lipid profile than humans. In humans, most circulating cholesterol (about 75%) is contained in very low density lipoprotein (V-LDL), whereas in mice, the level of V-LDL is very low, and most circulating cholesterol (about 80%) is contained in high density lipoprotein (HDL) [4]. Therefore, there are no inbred mouse strains that naturally develop atherosclerosis, which makes it challenging to establish models of atherosclerosis. To address this issue, studies have sought to establish models through genetic modification, and in 1992, the first transgenic model mouse, the Apolipoprotein E knockout (ApoE-/-) mouse, was reported through homologous recombination in embryonic stem cells [5]. ApoE is a protein that is synthesized mainly in the liver and brain. ApoE functions as a major ligand for the uptake of triglyceride-rich lipoprotein remnants in circulation into hepatocytes via low-density lipoprotein receptor, low-density lipoprotein receptor related protein 1, and heparan sulfate proteoglycans on the cell surface [6,7]. Because ApoE plays an important role in the clearance of lipoprotein remnants, ApoE−/− mice exhibit hyperlipidemia even when fed a normal diet [8,9], and this condition is exacerbated when they are given a high-fat or high-cholesterol diet [5, 10∼12]. After being fed a high-fat diet, ApoE-/- mice rapidly develop extensive atherosclerosis [13]. Plaque formation in ApoE-/- mice is very similar to that in established large animal models and humans, and many research groups continue to use this mouse model today [9]. However, ApoE-/- mice also have some limitations as a model for atherosclerosis. One of these limitations is that the most abundant lipoprotein in ApoE-/- mice is V-LDL, rather than low-density lipoprotein, which is characteristic of human atherosclerosis [5]. Additionally, in this model, plaque rupture and thrombosis, which cause human stroke and myocardial infarction, are rarely observed [14, 15]. Furthermore, human atherosclerosis is multifactorial and influenced by complex interactions, making it impossible to replicate all of its elements. To address these limitations, genetic modification alone cannot fully replicate the complex nature of human atherosclerosis. Recently, with continuing urbanization, obesity rates have increased, and various models of metabolic syndrome (METS) have been developed, many of which involve genetic modification. Establishing a mouse model of atherosclerosis associated with metabolic syndrome should better reflect current conditions. METS is a multifactorial metabolic disorder characterized by a cluster of metabolic abnormalities, including central obesity, dyslipidemia, hypertension, insulin resistance, glucose intolerance, and fasting hyperglycemia. A diagnosis of METS requires the presence of three of these five components [16]. At the root of METS are obesity and overnutrition, making it a representative lifestyle disease [17]. Additionally, METS doubles the risk for atherosclerotic cardiovascular disease [18]. Cardiovascular diseases are a leading cause of morbidity and mortality worldwide, and their incidence is expected to increase further by 2030 [19]. However, the mechanisms underlying the progression of atherosclerotic complications in METS are not yet fully understood, necessitating further elucidation. METS is a multifactorial disease with complex interactions, and all its components need to be studied together. The aim of this study was to create an animal model that more closely resembles the human pathophysiology by simultaneously reproducing hypertension, dyslipidemia, and glucose intolerance, which are components of METS.

## 4. Methods

### 4.1 Experimental protocol

All experimental protocols were approved by the Animal Care and Use Committee of Fukuoka University and conformed to the guide for the Care and use of laboratory animals of the Institute of Laboratory Animal Resources. ApoE-/- mice were purchased from Charles River Laboratories Japan, Inc., Japan. Twelve-week-old male ApoE-/- mice were started on a high-fat diet (HFD) for 12 weeks to create a condition of dyslipidemia. The high-fat diet (Clea Japan HFD32, CLEA Japan, Inc., Japan) contained 32g of fat. The proportions of calories from protein, fat, and nitrogen-free extract were 20%, 56.7%, and 23.1%, respectively. From 12 weeks, continuous subcutaneous administration of angiotensin II (ATII, 0.7 mg/kg per day) using an osmotic pump (ALZET osmotic pump, Muromachi Kikai Co., Ltd., Japan) was used for 12 weeks to create a hypertensive condition. The osmotic pump was replaced at 18 weeks based on its duration. At 18 weeks, streptozotocin (STZ) 80 mg/kg was administered intraperitoneally to create the condition of type 2 diabetes. By these means, we created a metabolic syndrome group (METS) that reproduced diabetes, hypertension, and dyslipidemia, an ATII group that received only angiotensin II, an STZ group that received only streptozotocin, and an HFD group that was fed only a high-fat diet. In addition, a control group (CTL) received no drugs and was fed a normal diet.

### 4.2 Body weight, blood pressure, and blood glucose

Body weight and blood pressure were measured every 4 weeks (weeks 12, 16, 20, 24). Blood pressure was measured through a tail cuff-based MK-2000 device (Muromachi Kikai Co., Ltd., Tokyo, Japan). Casual blood glucose was measured every 2 weeks from 18 weeks.

### 4.3 Biochemical examination

We sacrificed the mice at 24 weeks and collected their blood. We submitted the blood samples for analysis, including measurements of aspartate aminotransferase (AST), alanine aminotransferase (ALT), lactate dehydrogenase (LDH), blood urea nitrogen (BUN), creatinine (CRE), total cholesterol (T-CHO), triglycerides (TG), low-density lipoprotein cholesterol (LDL-C), high-density lipoprotein cholesterol (HDL-C), and hemoglobin A1c (HbA1c).

### 4.4 Echocardiography

Echocardiography was performed using isoflurane at weeks 12 and 24. Echocardiographic measurements were performed using a NEMIO SSA-550A (Toshiba, Tokyo, Japan). From the short-axis two-dimensional view and M mode at the level of the papillary muscle, we measured heart rate, interventricular septum thickness diameter (IVSd), left ventricular internal dimension in diastole (LVIDd), left ventricular posterior wall thickness diameter (LVPWd), left ventricular internal dimension in systole (LVIDs), left ventricular ejection fraction (LVEF), and left ventricular fractional shortening (LVFS).

### 4.5 Pathological analysis

The rate of atherosclerosis formation at 24 weeks was evaluated pathologically. The excised aorta was stained with Oil Red O stain and evaluated by the rate of atherosclerosis formation from the aortic origin to the renal artery bifurcation.

### 4.6 Histological analysis

We extracted ribonucleic acid (RNA) from the whole excised aorta by using a RiboPure RNA Purification Kit (LifeTechnologies, Carlsbad, CA, USA) and synthesized complementary Deoxyribonucleic acid (cDNA) by using a ReverTra Ace® qPCR RT Kit (TOYOBO, Japan). We lists the primers in Supplementary table. Reverse transcription-polymerase chain reaction (RT-PCR) of the synthesized cDNA was performed on a 7500 Fast Real-Time PCR System (Applied Biosystems, USA) using THUNDERBIRD® SYBR® qPCR Mix (TOYOBO, Japan). We measured receptors and ligands that are expressed on vascular endothelial cells and have been specifically implicated in atherosclerosis, such as Nicotinamide Adenine Dinucleotide Phosphate Oxidase2 (NOX2), Nuclear Factor Kappa B1, Matrix Metalloproteinase 2 (MMP-2), Integrin Subunit Alpha 2b (Itga2b), vascular endothelial growth factor receptor-2 (VEGFR-2), β-cadherin-related proteins (β-catenin), nuclear factor erythroid 2-related factor 2 (Nrf2), and Heme Oxygenasse-1 (HO-1). Results were analyzed with β actin as a housekeeping gene.

### 4.7 Statistical analysis

All data analyses were performed using SAS (version 9.4, SAS Institute Inc., Cary, NC, USA) at Fukuoka University (Fukuoka, Japan). Continuous variables were expressed as mean ± standard deviation. The five groups, CTL, HT, ATII, STZ, HFD, and METS groups, were evaluated by analysis of variance, with a p-value of <0.05 indicating statistical significance.

## 5. Results

### 5.1 Changes in body weight

The changes in body weight are summarized in Figure 1. At 12 weeks of age, the baseline body weight was 27.8 ± 1.5 g. At 24 weeks of age, the average body weight was 34.6 ± 5.4 g. After 16 weeks of age, the HFD and METS groups showed a significant increase in body weight compared to the other groups.

**Figure 1.**
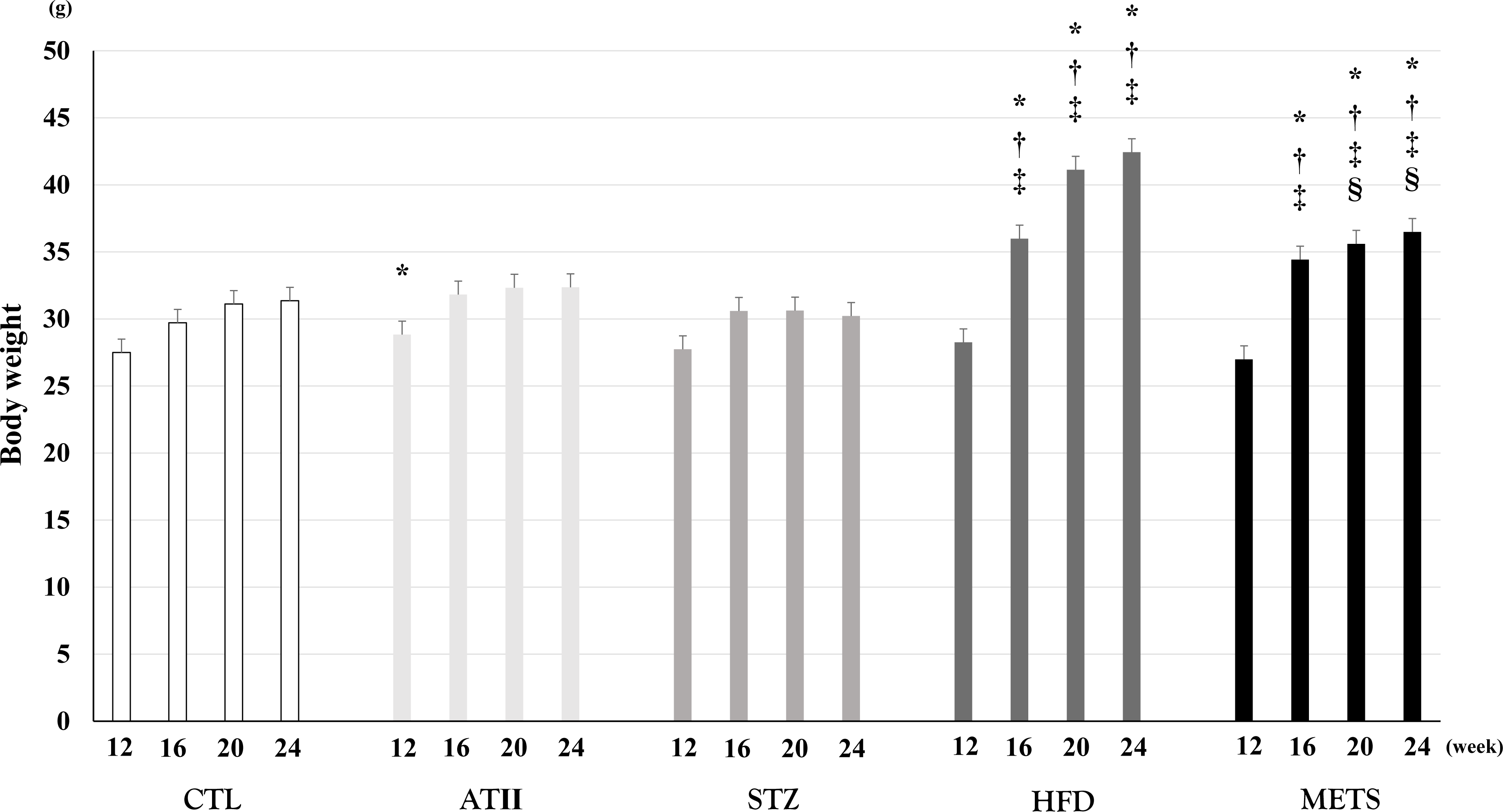
Changes in body weight. Changes in body weight in each group. CTL, control group; ATII, AngiotensinII group; STZ, streptozotocin group; HFD, high-fat diet group; METS, the group administered angiotensinII, streptozotocin, and a high-fat diet. CTL (n = 17), ATII (n = 18), STZ (n = 18), HFD (n = 17), and METS (n = 21) were investigated. *, †, ‡, and § indicate significant differences compared with CTL, ATII, STZ, and HFD during the same week, respectively.

### 5.2 Changes in blood pressure

The changes in blood pressure are summarized in Figure 2. At 12 weeks of age, the mean systolic blood pressure was 100.4 ± 14.1 mmHg. At 24 weeks of age, the mean systolic blood pressure was 117.6 ± 22.9 mmHg. At 16 and 20 weeks, the STZ group showed a significant decrease in systolic blood pressure.

**Figure 2.**
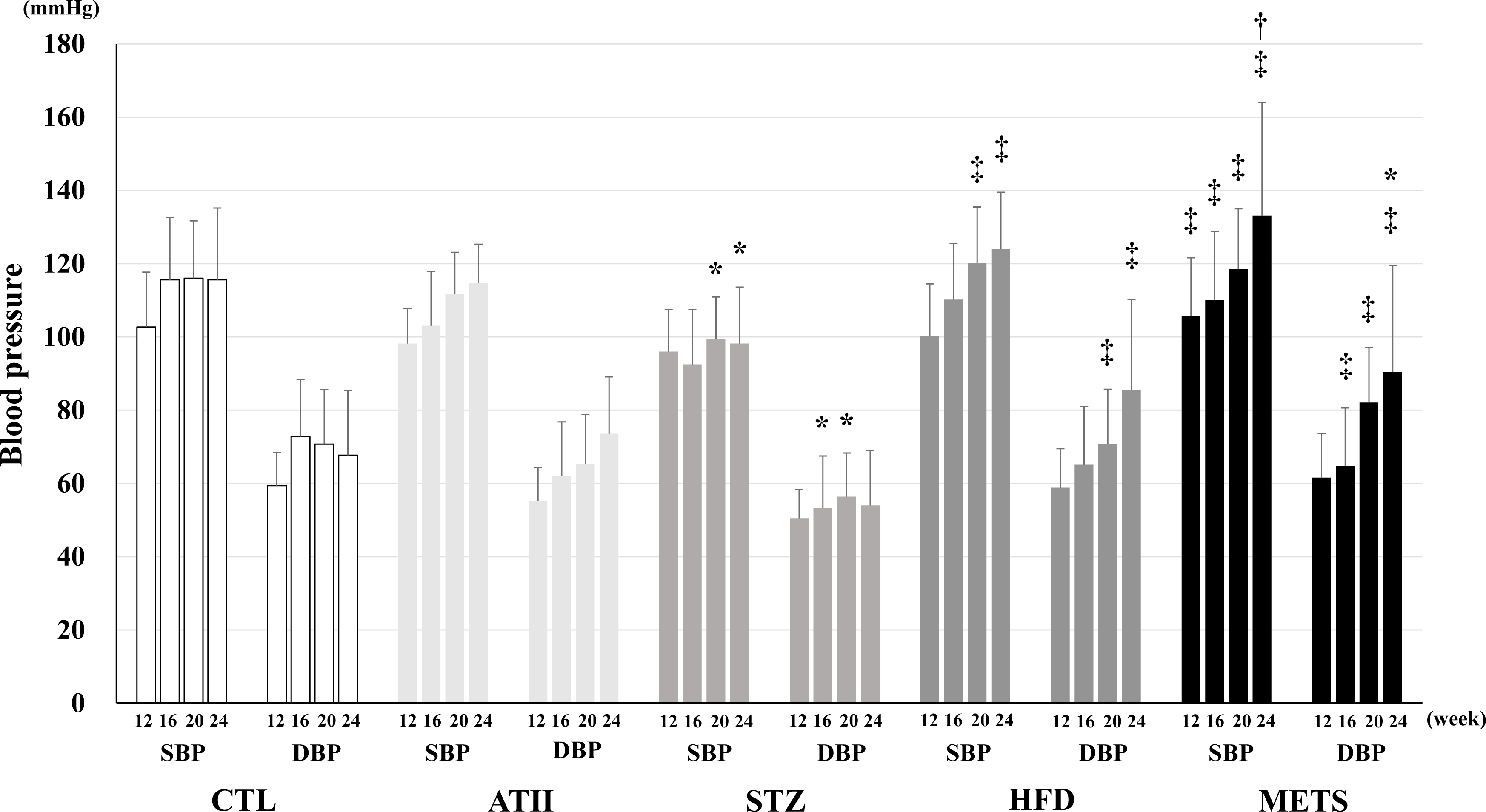
Changes in blood pressure. Changes in blood pressure including systolic blood pressure and diastolic blood pressure in each group. CTL, control group; ATII, AngiotensinII group; STZ, streptozotocin group; HFD, high-fat diet group; METS, the group administered angiotensinII, streptozotocin, and a high-fat diet. CTL (n = 17), ATII (n = 18), STZ (n = 18), HFD(n = 17), and METS(n = 21) were investigated. *, †, ‡, and § indicate significant differences compared with CTL, ATII, STZ, and HFD during the same week, respectively.

### 5.3 Blood examinations

The results of blood examinations at 24 weeks are shown in Table 1. In the HFD group, liver enzymes such as AST, ALT, and LDH were significantly elevated. Regarding kidney function, BUN was significantly lower in the HFD group, while a significant decrease in CRE was observed in the METS group. For the lipid profile, LDL-C increased in the METS group, but a significant difference was observed only compared to the ATII group. Additionally, HDL-C was significantly higher in the METS group compared to the other groups. There were no significant differences in HbA1c among the groups. Additionally, we observed the blood glucose trends at 18, 20, 22, and 24 weeks in each group (Figure 3). At 18 weeks of age, the mean blood glucose was 215.3±42.2 mg/dL. At 24 weeks of age, the mean blood glucose was 220±59.3 mg/dL. There were no significant differences between the groups.

**Figure 3.**
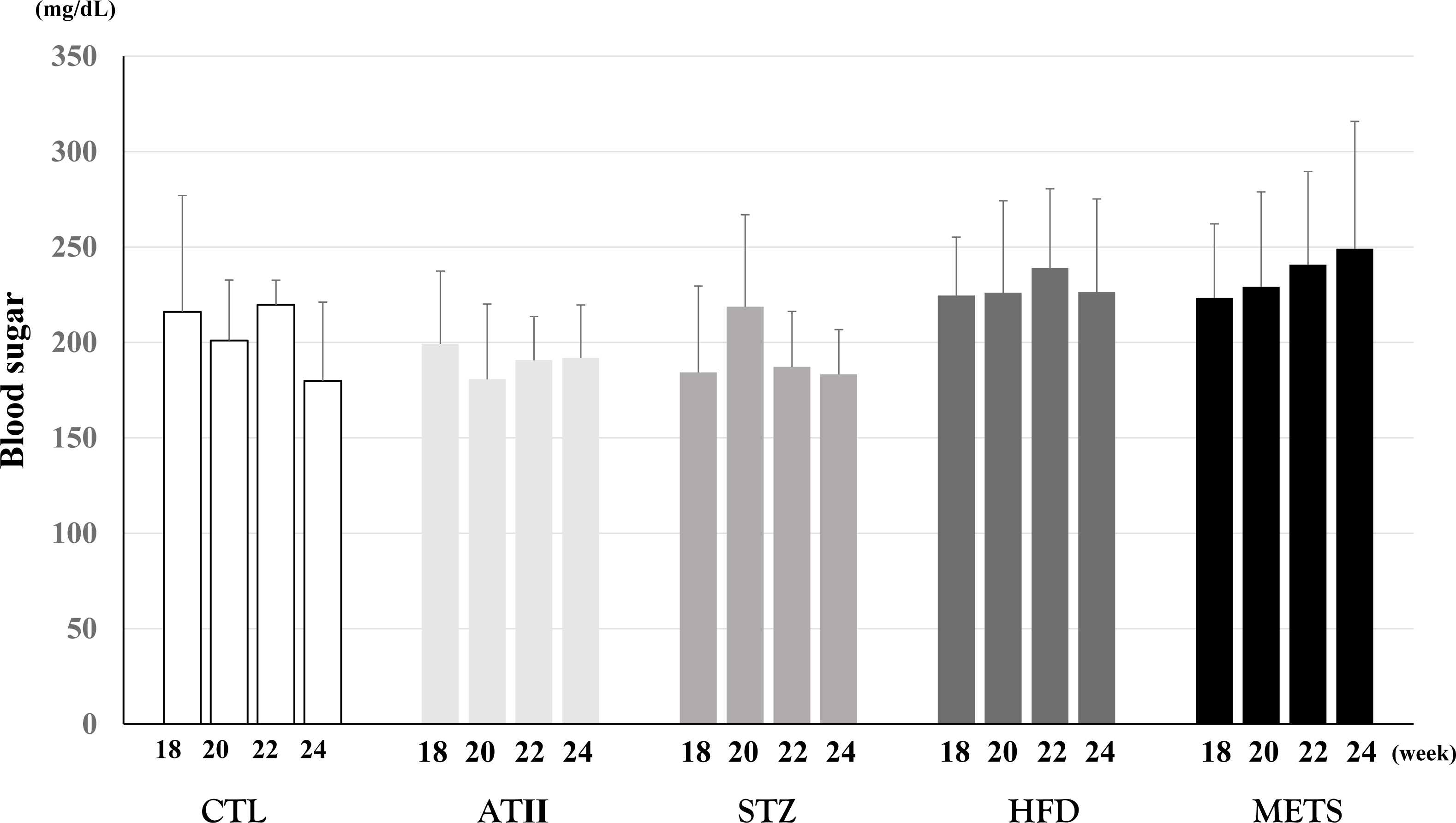
Changes in blood sugar. Changes in blood sugar in each group. CTL, control group; ATII, Angiotensin group; STZ, streptozotocin group; HFD, high-fat diet group; METS, the group administered angiotensin, streptozotocin, and a high-fat diet. The round marker and solid line, square marker and dotted line, triangle marker and dashed line, rhombus marker and chain line, and cross marker and double line indicate CTL, ATII, STZ, HFD, and METS, respectively. CTL (n = 17), ATII (n = 18), STZ (n = 18), HFD(n = 17), and METS(n = 21) were investigated. *, †, ‡, and § indicate significant differences compared with CTL, ATII, STZ, and HFD during the same week, respectively.

**Table 1.** Blood test at 24 weeks.

|  | CTL | ATII | STZ | HFD | METS |
| --- | --- | --- | --- | --- | --- |
| AST, IU/L | 67.8±36 | 48.7±8.7 | 39.5±8.6 | 467.2±250 *†‡ | 99.7±46.6 § |
| ALT, IU/L | 19.8±5.34 | 21±4.9 | 18.3±1.96 | 318.5±96.2 *†‡ | 80.8±59.5 § |
| LDH, IU/L | 220±82 | 179.7±26 | 131.5±21.6 | 1068±695 *†‡ | 480.7±139 § |
| BUN, mg/dL | 39.05±6.7 | 36.9±2.08 | 37.7±2.21 | 26.4±2.35 *†‡ | 42.6±9.07 § |
| CRE, mg/dL | 0.14±0.026 | 0.14±0.06 | 0.092±0.05 | 0.098±0.02 | 0.06±0.016 *† |
| T-CHO, mg/dL | 669.5±209 | 559.8±43.8 | 735.3±90.3 | 506.7±101.5 ‡ | 560±85.1 |
| TG, mg/dL | 84.7±36.7 | 74.8±17.9 | 106±56.3 | 100.8±29.3 | 150.6±56.8 |
| LDL-C, mg/dL | 112.8±38.2 | 88.2±22.1 | 119±12.8 | 115.8±13.9 | 146.8±26.4 † |
| HDL-C, mg/dL | 10.7±3.9 | 14.7±3.7 | 14.3±3.3 | 21.5±7.7 * | 34.6±2.7 *†‡§ |
| HbA1c, % | 5±0.2 | 4.8±0.15 | 4.9±0.25 | 4.9±0.32 | 5±0.32 |

CTL: control group, ATII: angiotensin II group, STZ: streptozotocin group, HFD; high fat
diet group, METS: the group which administrated angiotensin2, streptozotocin, and HFD,
AST: aspartate aminotransferase, ALT: alanine aminotransferase, LDH: lactate
dehydrogenase, BUN: blood urea nitrogen, CRE: creatinine, T-CHO: total cholesterol, TG:
triglycerides, LDL-C: low-density lipoprotein cholesterol, HDL-C: high-density lipoprotein
cholesterol, HbA1c: hemoglobin A1c. \*, †, ‡, and § show significant differences compared to CTL, ATII, STZ, and HFD, respectively.

### 5.4 Cardiac function and heart weight

We investigated cardiac function using echocardiography at 12 and 24 weeks of age. At 12 weeks of age, the mean LVIDd was 4±0.25 mm, the mean LVIDs was 2.28±0.27 mm, and the mean LVEF was 80±5.3 %. At 24 weeks of age, the mean LVIDd was 4.1±0.3 mm, the mean LVIDs was 2.4±0.36 mm, and the mean LVEF was 77±6.7%. Figure 4A-G shows the difference between the echocardiography parameters at 24 weeks and 12 weeks. There were no significant differences in the changes in these parameters over the 12-week period. We also examined the weights of hearts excised at 24 weeks (Figure 5). The mean heart weight was 160±24 mg. There were no significant differences between the groups.

**Figure 4.**
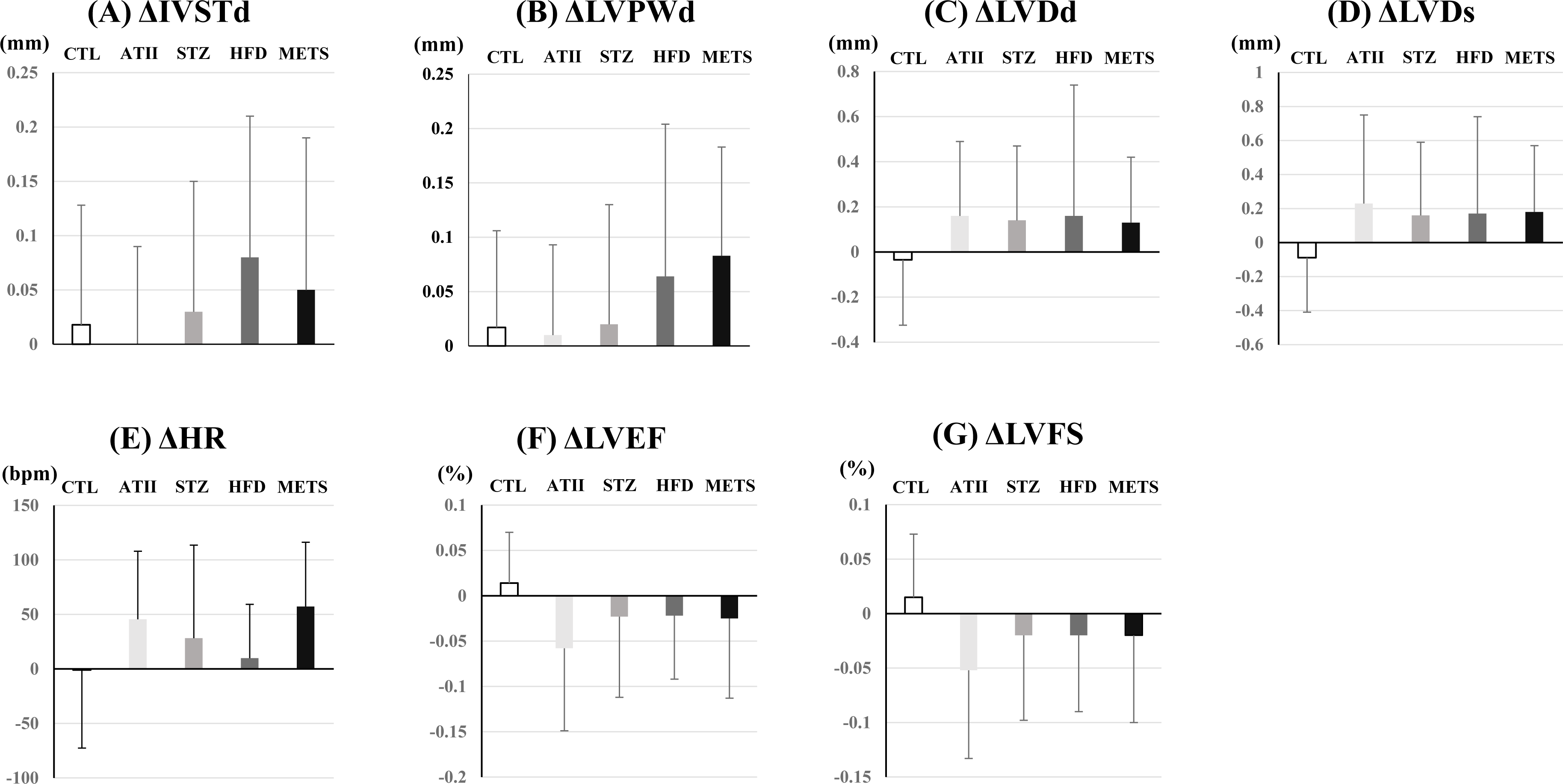
Changes in cardiac parameters via echocardiography after 12 weeks of treatment. Changes in IVSd (A), LVPWd (B), LVIDd (C), LVIDs (D), HR (E), LVEF (F), and LVFS (G) in each group. IVSd, interventricular septum thickness diameter; LVPWD, left ventricular posterior wall thickness diameter; LVIDd, left ventricular internal dimension in diastole; LVIDs, left ventricular internal dimension in systole; HR, heart rate; LVEF, left ventricular ejection fraction; LVFS, left ventricular fractional shortening; CTL, control group; ATII, Angiotensin group; STZ, streptozotocin group; HFD, high-fat diet group; METS, the group administered angiotensin, streptozotocin, and a high-fat diet. CTL (n = 17), ATII (n = 18), STZ (n = 18), HFD(n = 11), and METS (n = 12) were investigated. No significant difference was observed between the groups in this figure.

**Figure 5.**
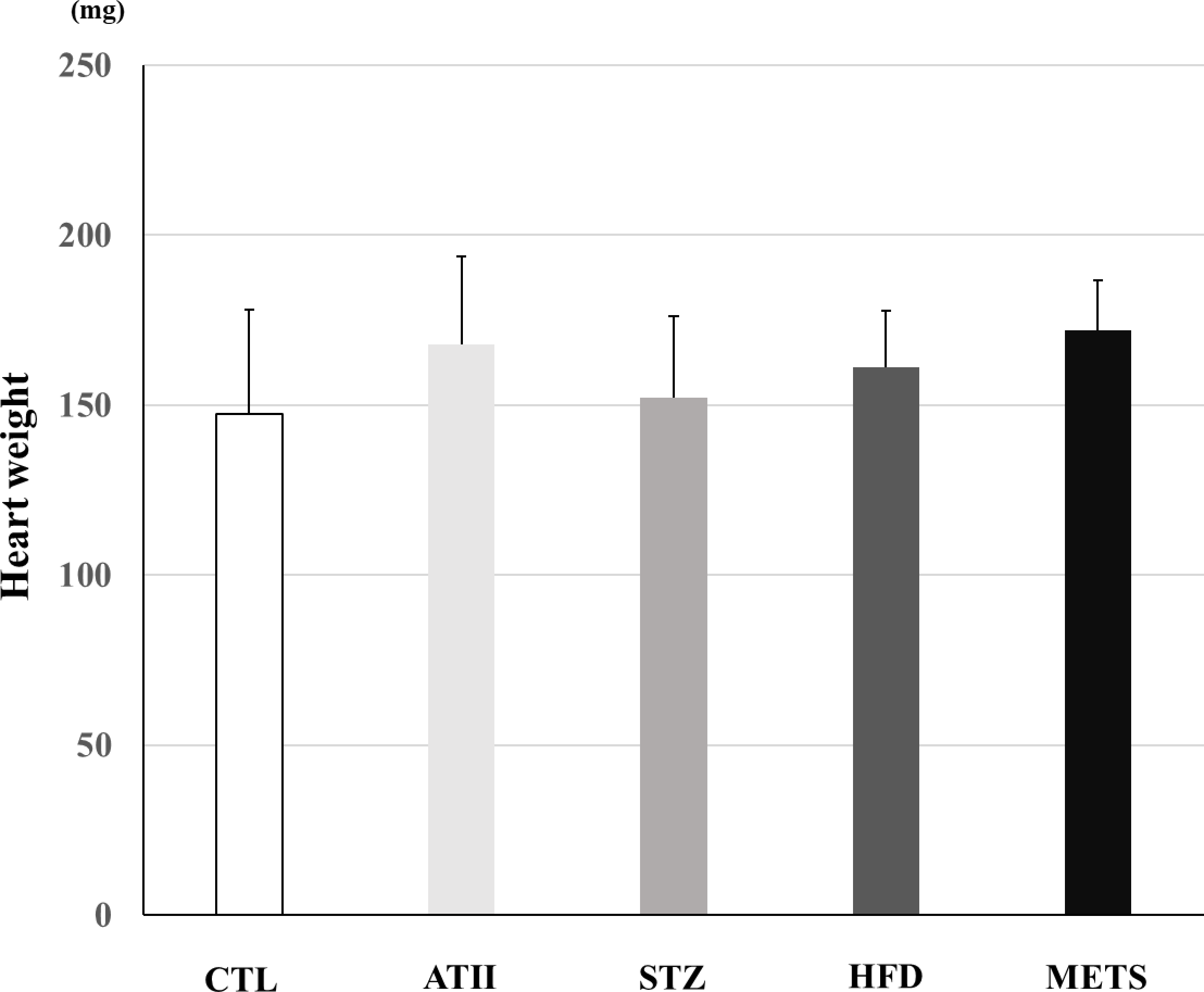
Heart weight. Weights of heart excised at 24 weeks in each group. CTL, control group; AtII, AngiotensinII group; STZ, streptozotocin group; HFD, high-fat diet group; METS, the group administered angiotensinII, streptozotocin, and a high-fat diet. CTL (n = 12), AtII (n = 12), STZ (n = 12), HFD(n = 11), and METS(n = 11) were investigated. *, †, ‡, and § indicate significant differences compared with CTL, AtII, STZ, and HFD, respectively.

### 5.5 Arteriosclerosis formation

We stained the excised arteries with Oil Red O at 24 weeks. Representative samples and the percentage of the area stained with Oil Red O for each group are shown in Figure 6. The METS group showed a significantly higher rate of atherosclerosis formation, followed by the ATII group. In the other groups, atherosclerosis formation was minimal.

**Figure 6.**
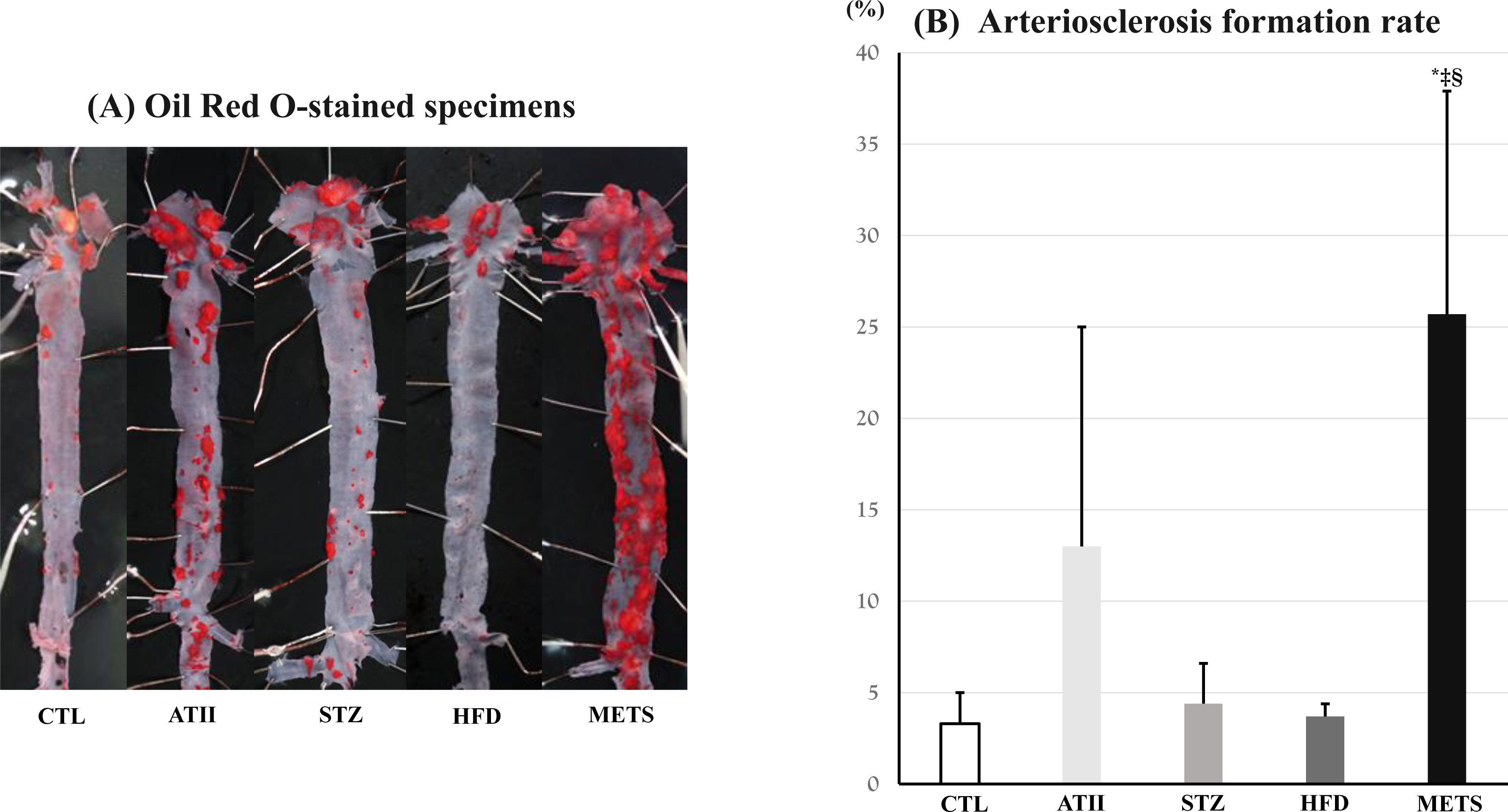
Arteriosclerosis formation. Oil Red O-stained specimens(A) and the arteriosclerosis formation rate (B) in each group. CTL, control group; AtII, Angiotensin group; STZ, streptozotocin group; HFD, high-fat diet group; METS, the group administered angiotensin, streptozotocin, and a high-fat diet. CTL (n = 5), AtII (n = 6), STZ (n = 6), HFD(n = 6), and METS(n = 10) were investigated. *, †, ‡, and § indicate significant differences compared with CTL, AtII, STZ, and HFD, respectively.

### 5.6 Advanced glycation end products (AGEs) receptor for the AGEs (RAGE) signaling pathway in complications of diabetes

We investigated messenger RNA (mRNA) expression in the AGEs RAGE signaling pathway, a biochemical pathway that plays a crucial role in the onset and progression of diabetic complications (Figures 7A-C). As a result, the METS group showed significantly higher expression levels of NOX2, Nuclear Factor Kappa B1, and MMP-2 compared to the CTL and HFD groups.

**Figure 7.**
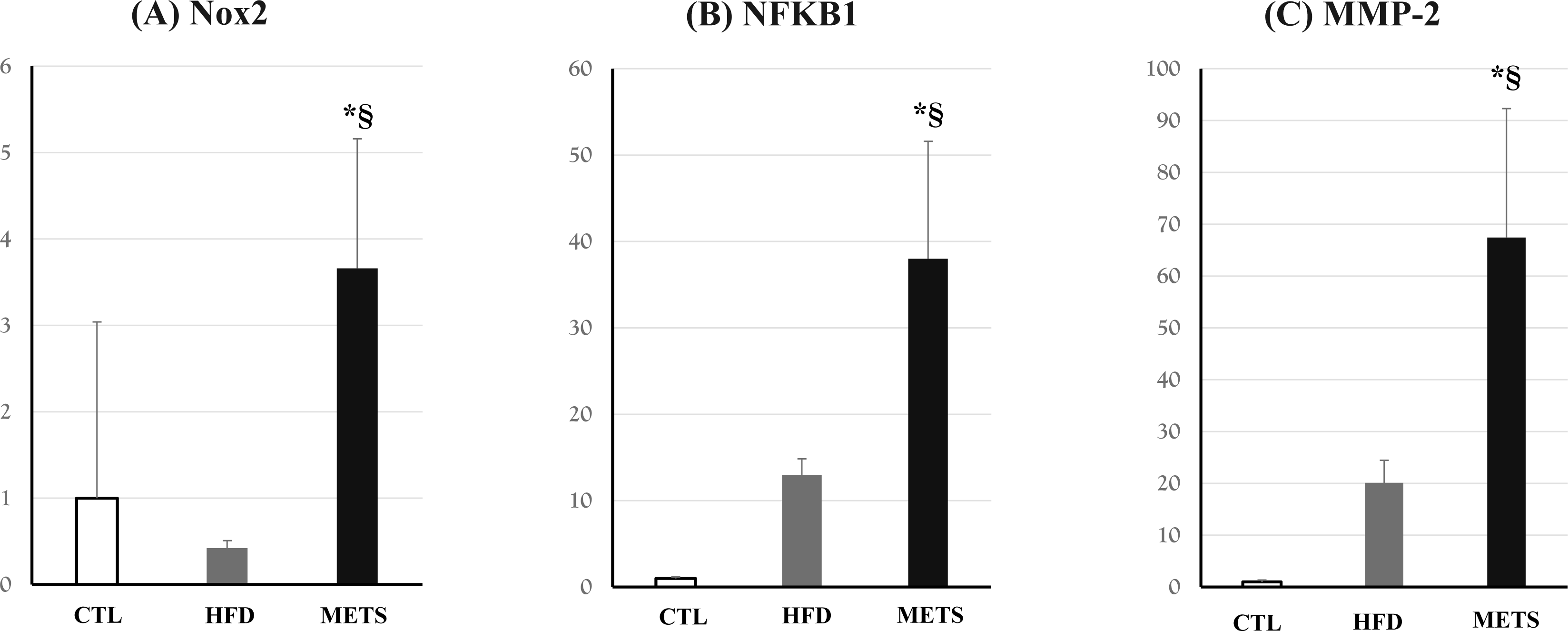
Age-RAGE signaling pathway in complications of diabetes. NOX2, nicotinamide adenine dinucleotide phosphate oxidase 2 (A); NFKB1, nuclear factor-kappa B1 (B); MMP-2, matrix metalloproteinase-2 (C) in each group. CTL, control group; HFD, high-fat diet group; METS, the group administered angiotensin, streptozotocin, and a high-fat diet. CTL (n = 5), HFD(n = 5), and METS(n = 5) were investigated.§indicates a significant difference compared with HFD.

### 5.7 Fluid shear stress and atherosclerosis

We investigated mRNA expression during the progression of atherosclerosis induced by a fluid stress. These pathways regulate the defensive response against atherosclerosis (Figures 8A-E). As a result, the METS group showed significantly higher expression levels of Itga2b, Nrf2, and HO-1 compared to the CTL group.

**Figure 8.**
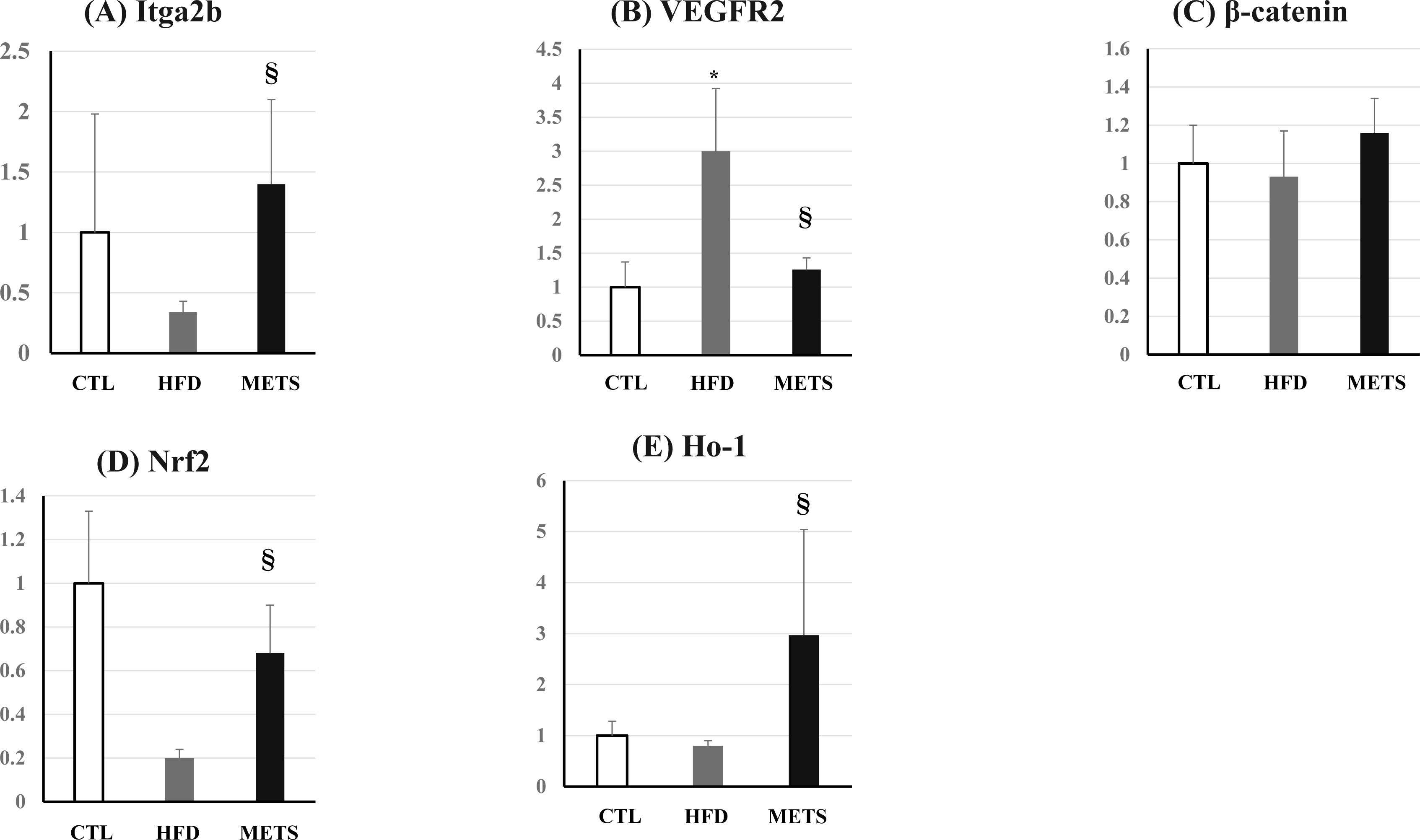
Fluid shear stress and atherosclerosis. Itga2b, Integrin Subunit Alpha 2b (A); VEGFR-2, vascular endothelial growth factor receptor-2 (B); β-catenin, β-cadherin-related proteins(C); Nrf2, nuclear factor erythroid 2-related factor 2 (D); HO-1, Heme oxygenase-1 (E) in each group. CTL, control group; HFD, high-fat diet group; METS, the group administered angiotensin, streptozotocin, and a high-fat diet. CTL (n = 5), HFD(n = 5), and METS(n = 5) were investigated. *§ indicate significant differences compared with CTL and HFD, respectively.

## 6. Discussion

### 6.1 Arteriosclerosis

In this study, we confirmed significantly higher arteriosclerosis formation rates in the METS group (Figure 6). There was only slight arteriosclerosis in the CTL, STZ, and HFD groups. The ATII group exhibited a noticeable amount of arteriosclerosis formation. These results suggest that, in addition to the atherosclerosis induced by ATII, the load of STZ or HFD further accelerates atherosclerosis formation. The mRNA expression levels of NOX2 and Nuclear Factor kappa-light-chain-enhancer of activated B cells (NF-κB) were significantly elevated in the METS group (Figure 7), suggesting the influence of oxidative stress. The oxidative stress induced by increased reactive oxygen species can potentially damage various intracellular structures such as cell membranes, proteins, lipids, and DNA. Reactive oxygen species products include hydrogen peroxide, superoxide anion, hydroxyl radical, and nitric oxide, which are generated by mitochondrial oxidase, NOX, and nitric oxide synthase [20]. Additionally, a significant increase in the mRNA expression of MMP-2 was observed in the METS group. MMP-2 activates the NF-κB pathway, promoting the expression of inflammatory cytokines and advancing atherosclerosis [21]. Furthermore, MMP-2 is known to promote remodeling of the extracellular matrix through the Transforming Growth factor-β signaling pathway [22]. In this study, we believe that the METS load successfully replicated the increase in oxidative stress and inflammatory cytokines associated with diabetes-induced atherosclerosis formation. We also investigated mediators related to shear stress and atherosclerosis. Endothelial cells (ECs) are constantly exposed to shear stress due to the frictional forces generated by blood flow. In response to shear stress, ECs regulate intracellular signaling, leading to changes in gene expression, cell morphology, and structural remodeling [23,24]. Actually, the endothelium in the curved regions of the aorta and atherosclerotic areas shows increased signaling expression and activation compared to non-atherosclerotic areas and the straight parts of the aorta. The application of shear stress to ECs is known to activate multiple mechanosensors on the cell membrane. These include Itga2b and VEGFR-2. Activation of VEGFR-2 is associated with β-catenins. In this study, the mRNA expression of Itga2b was significantly elevated in the METS group, whereas the mRNA expression of VEGFR-2 did not increase in the METS group but was elevated in the HFD group (Figure 8). There was no difference in β-catenin among the groups. The activation of integrins leads to the activation of mitogen-activated protein kinase. The activation of mitogen-activated protein kinase contributes to cell proliferation and differentiation in response to stress. VEGFR-2 and β-catenin phosphorylate downstream protein kinase B [25]. The activation of β-catenin phosphorylate downstream protein kinase B contributes to the survival of endothelial cells and the suppression of inflammatory responses, thereby playing a role in anti-atherosclerosis [26,27]. In this study, the mRNA expression levels of Nrf2 and HO-1 were measured as downstream mediators related to shear stress and atherosclerosis. Nrf2 was not increased compared to that in the CTL group, but HO-1 was significantly elevated. Nrf2 is a transcription factor that regulates antioxidant responses and protects cells from oxidative stress. Shear stress promotes the nuclear translocation of Nrf2 [28], increasing the expression of antioxidant enzymes such as HO1, which plays a crucial role in the prevention of atherosclerosis. Additionally, Nrf2 not only activates the expression of many antioxidant enzymes but also has the potential to regulate NOX activity and may be associated with the Age-RAGE pathway. In most normal tissues, HO1 is insufficiently expressed, but its expression increases when vascular injury occurs [29]. Additionally, HO1 is highly expressed in atherosclerotic lesions and correlates with plaque instability and pro-inflammatory markers [30]. The significant increase in HO-1 in the METS group is considered to reflect an increase in shear stress and an enhanced defensive response against atherosclerosis formation.

### 6.2 Hypertension

Activation of the renin-aldosterone-angiotensin system is a common feature in patients with metabolic syndrome, who have a fourfold higher risk of developing atherosclerosis [31–33]. The ATII system is involved in the development of atherosclerosis, which not only leads to the production of reactive oxygen species but also to an increase in the expression of inflammatory gene products. Thus, one of the most widely used preclinical models of hypertension, particularly in rodents, involves the long-term subcutaneous infusion of ATII [34]. The American Heart Association suggests the following doses of ATII for creating hypertensive mouse models [35]: slow pressor dose (5.76 mg/kg/day) [36], intermediate dose (7.05-7.2 mg/kg/day) [37], and higher dose (14.4 mg/kg/day). A slow pressor dose may mimic the gradual onset of hypertension in humans with primary hypertension. An intermediate dose has been the most widely used in recent years. ATII induces abdominal aortic aneurism when administered at high doses [38]. In this experiment, we focused on atherosclerosis, and therefore, the occurrence of abdominal aortic aneurism was undesirable, leading us to use the intermediate dose. As a result, no groups in this study exhibited an average systolic blood pressure exceeding 140 mmHg, and only the METS group showed a diastolic blood pressure exceeding 90 mmHg at 24 weeks (Figure 2). The highest systolic blood pressure at 24 weeks was observed in the METS group, followed by the HDL group. No significant increase in blood pressure was observed in the ATII group.

### 6.3 Dyslipidemia

In this study, significant weight gain was observed in the HFD and METS groups, with the HFD group gaining more weight than the METS group (Figure 1). The STZ group had the lowest weight trajectory, but there was no significant difference. The difference in weight between the HFD group and the METS group is thought to be influenced by the administration of STZ to the METS group. In fact, many previous studies have reported weight loss in mice administered low doses of STZ [39]. Blood tests conducted at 24 weeks showed no differences in total cholesterol levels between the groups (Table 1). An increase in LDL-C and HDL-C was observed in the METS group, but not in the HFD group. In this experiment, all of the groups consisted of ApoE -/- mice, which are inherently prone to hyperlipidemia. However, as mentioned earlier, it is known that the administration of a high-fat diet to ApoE-/- mice results in further hypercholesterolemia, which was not observed in the HFD group in this study. The changes in the lipid profile observed in the METS group are thought to be influenced by insulin resistance and hypertension. Additionally, liver function abnormalities were observed in both the HFD and METS groups, with the HFD group showing more pronounced changes. Generally, the creation of non-alcoholic fatty liver disease (NAFLD) model mice involves the administration of a Western diet containing more than 45% of calories from fat [40]. In this experiment, we administered a Western diet containing 56.7% of calories from fat, which is believed to induce NAFLD through high-calorie intake and increased adiposity. NAFLD is typically associated with obesity [41], and the differences in liver function abnormalities between the HFD and METS groups are considered to be due to their differences in body weight.

### 6.4 Diabetes Mellitus

Diabetes Mellitus (DM) is ordinarily divided into type1 DM (T1DM), which is caused by an absolute insulin deficiency, and type2 DM (T2DM), which is caused by a relative insulin deficiency. Dietary manipulations such as high-fat diets and fructose-rich diets have been used to reproduce DM, but these models require a long time to induce diabetes. Therefore, a faster model using STZ has been developed, and since its discovery in 1959, it has been widely used to induce diabetes in experimental animals and preclinical trials [42,43]. STZ is an alkylating antitumor agent. Because it is a glucose analog, it enters β-cells via the glucose transporter type2 and accumulates intracellularly. Intracellular STZ forms diazomethane, an alkylating agent, which causes DNA methylation and induces diabetic effects [44]. STZ can reproduce T1DM at high doses and T2 DM due to the partial destruction of beta cells at low doses. It is also known that feeding a high-fat diet in addition to low-dose STZ administration can increase insulin resistance, making the model more similar to human T2 DM [45]. After STZ administration, mice exhibit increased food intake, decreased mobility, and behavioral changes such as piloerection [46]. In this study, we aimed to create a state of type 2 diabetes by administering low-dose STZ from 18 weeks. For the first four weeks after administration, there was no significant difference in random blood glucose levels among the groups. However, at the sixth week, a significant increase in blood glucose was observed in the METS group, exceeding the generally considered hyperglycemic level of 250 mg/dL in mice (Figure 3). No increase in blood glucose was observed in the STZ group, supporting the idea that not only pancreatic weakening but also conditions like hypertension and hyperlipidemia bring the model closer to actual diabetes in humans.

In conclusion, the METS model mouse showed severe atherosclerosis caused by oxidative stress, inflammation, and atherosclerotic plaque formation. This model is expected to be useful as a model that more closely resembles the pathophysiology of clinical atherosclerosis.

## Non-standard Abbreviations and Acronyms

METS: metabolic syndrome
CTL: Control
HFD: high-fat diet
ATII: streptozotocin
STZ: angiotensin II
NOX: Nicotinamide Adenine Dinucleotide Phosphate oxidase
MMP-2: Matrix Metalloproteinase 2
VEGFR-2: vascular endothelial growth factor receptor-2
β-catenin: β-cadherin-related proteins
Nrf2: nuclear factor erythroid 2-related factor 2
HO-1: Heme Oxygenasse-1
AGEs: Advanced glycation end products
RAGE: receptor for the AGEs

## Acknowledgements

We thank Kanji Nakai, Sayo Tomita, and Satomi Abe for their excellent technical assistance.

